# Plasticity of transmembrane helix interactions in EphA2 dimers and oligomers

**DOI:** 10.1101/2022.06.06.495010

**Authors:** Daniel Wirth, Ece Ozdemir, William C. Wimley, Elena B. Pasquale, Kalina Hristova

## Abstract

Lateral interactions can stabilize different EphA2 receptor assemblies in the plasma membrane in response to different ligands. Here we use two fluorescent techniques, Forster Resonance Energy Transfer (FRET) and Fluorescence Intensity Fluctuations (FIF) spectrometry, to investigate how mutations in the EphA2 transmembrane (TM) helix affect the association between full-length EphA2 molecules in the absence of ligand and in the presence of three ligands: ephrinA1-Fc, m-ephrinA1, and the YSA peptide. The EphA2 mutations we studied have been previously characterized in the context of the isolated EphA2 TM helix. Working with full-length EphA2, we observed modest effects of the mutations on receptor-receptor interaction. Our data do not support the currently accepted model of a switch between two discrete TM helix dimerization motifs corresponding to active or inactive receptor states. Instead, we propose that different dimeric/oligomeric arrangements of the EphA2 extracellular region couple to an ensemble of TM helix dimer interfaces. Plasticity in the arrangements of receptor tyrosine kinase TM helices in active dimers and oligomers may serve to facilitate the cross-phosphorylation of multiple tyrosines in different positions of the intracellular regions.

## Introduction

Receptor tyrosine kinases (RTKs) are single-pass transmembrane proteins that control cell growth, differentiation, motility and metabolism (1-3). They transduce biochemical signals via lateral oligomerization in the plasma membrane. The catalytic activity of the intracellular kinase domains is stimulated by cross-phosphorylation of neighboring RTK molecules in dimers and oligomers, which results in the activation of downstream signaling cascades that control cell behavior (2,4-6). Dysregulation of RTK activity has been linked to many human diseases, including most cancers (2,3,7,8). Thus, RTKs are promising drug targets, and a number of RTK inhibitors are already used in the clinic (9-12).

The single transmembrane (TM) helix embedded in the plasma membrane connects the extracellular and intracellular portions of an RTK and may thus play a critical role in signal transduction across the plasma membrane. Work in the past two decades has demonstrated that contacts between TM helices can contribute to the overall stability of RTK dimers and oligomers (13-16). Furthermore, RTK TM helices have been proposed to interact via two specific dimerization motifs characteristic of either inactive or active RTK states, suggesting a role of the TM helix in RTK activation.(17-22). The switch from one well-defined TM dimer structure to the other, occurring upon ligand binding, could be a mechanism enabling transmission of information about the presence of a bound ligand to the kinases domain (23). However, this concept of a “TM dimer switch” has been mainly supported by experimental data obtained with isolated TM helices and by computational modeling (20-22,24). On the other hand, the consequences of TM helix mutagenesis on the function of full-length RTKs have been difficult to interpret based simply on the TM dimer switch model (25). Therefore, the role of TM helices in RTK signal transduction is still unclear, despite many years of research. Recent cryo-EM structures of RTKs have also not provided insights into the role of the TM helix in RTK signaling because the TM helices have remained unresolved in these structures (26,27).

We seek to understand the role of the TM helix in the activation of EphA2, an RTK that plays an important role in cancer, inflammation, atherosclerosis, and infections (28-30). The extracellular region of EphA2 (including an N-terminal ligand-binding domain, a cysteine-rich region, and two fibronectin type III domains) (31,32). The EphA2 TM helix is connected to the tyrosine kinase domain by a flexible juxtamembrane segment of ∼50 amino acids (31,32). The EphA2 intracellular portion also includes a SAM domain, after the kinase, and a short C-terminal tail. Interactions between EphA2 molecules in the plasma membrane are complex, since EphA2 can form not only dimers but also higher order oligomers (31-33). Oligomerized EphA2 molecules phosphorylate each other on tyrosines in the juxtamembrane segment, the kinase domain and the SAM domain (34). Tyrosine phosphorylation promotes EphA2 kinase activity and downstream signaling controlling cell morphology, adhesion, migration, proliferation, and survival (35-39). EphA2 kinase-dependent signaling in tumors can suppress the AKT-mTORC1 and RAS-ERK oncogenic pathways and inhibit cell adhesion and migration/invasion, but can also promote tumor angiogenesis and the dispersal of cancer cells (34).

The structure of the dimeric EphA2 TM helix, embedded in lipid bicelles mimicking the plasma membrane, has been solved by NMR (20). These studies utilized a peptide that includes EphA2 residues 523 to 563, encompassing a short N-terminal hydrophilic segment (corresponding to the end of the second fibronectin type III domain and an extracellular 7 amino acid linker), the hydrophobic membrane-embedded sequence of the TM helix, and the HRRRK stop-transfer sequence representing the positively charged N-terminal portion of the juxtamembrane segment. In the bicelles, the EphA2 TM helices interact via an extended “heptad repeat (HR)” motif that includes residues G539, A542 and G553. A shorter segment, comprising EphA2 residues 531 to 563 including the TM helix and the N-terminal portion of the juxtamembrane segment, dimerizes in cells via a different “glycine zipper (GZ)” interface involving residues G540 and G544 (24). These and other studies of the isolated EphA2 TM helix have been interpreted on the basis of the ligand-induced “TM dimer switch” model, which involves a switch in the conformation of dimerized TM helices from an inactive conformation to a different active conformation (19,21). However, the relevance of this model to the behavior of full-length EphA2 in the plasma membrane is unknown.

Functional studies of mutations, engineered to destabilize either the HR or the GZ interface in full-length EphA2, do not support the ligand-induced TM dimer switch model (19). Although mutations in the HR and GZ motifs increase and decrease EphA2 tyrosine phosphorylation, respectively, as compared to the wild-type receptor, the effects of the mutations are relatively modest and do not depend on ligand binding. Other recent findings further raise the possibility that the specific arrangement of the TM helices in EphA2 dimers and oligomers may not be critical for receptor activation. For instance, deletion of most of the juxtamembrane segment (residues 565 to 606) severely compromises the ability of EphA2 molecules to cross-phosphorylate on the remaining tyrosine residues (including Y772 in the activation loop of the kinase domain and Y930 in the SAM domain) (40). This suggests that the flexible juxtamembrane sequence, connecting the TM helix with the kinase domain, allows differential positioning of EphA2 molecules for cross-phosphorylation on different tyrosine residues. The flexibility of this segment argues against tight structural coupling between the TM helix and the kinase domain and thus against a major impact of TM dimeric arrangements on EphA2 tyrosine phosphorylation and activation.

EphA2 can be differentially activated by multiple ligands. The ligand most widely used to activate EphA2 is the dimeric ephrinA1-Fc (34,41), a chimeric protein composed of ephrinA1 fused to an antibody Fc region. EphrinA1-Fc potently promotes EphA2 kinase-dependent signaling (34,42). The endogenous form of ephrinA1 is anchored on the cell surface by a glycosylphosphatidylinositol (GPI) linkage, but can also be released in a monomeric form (m-ephrinA1) that can also activate EphA2 (40,43). We recently found that these are biased ligands (40). We also found that ligands do not always cause substantial EphA2 activation. For instance, the small monomeric peptide ligand YSA-GSGSK (YSAYPDSVPMMSGSGSK) activates EphA2 only very weakly (40,44). Although the three ligands can stabilize different EphA2 oligomeric assemblies in the plasma membrane of live cells (45), it is not known whether the TM helix plays a role in these assemblies.

Here we investigate how mutations in the HR and GZ interfaces of the EphA2 TM helix affect the dimerization/oligomerization of EphA2 molecules in cells in the absence of ligand and in the presence of ephrinA1-Fc, m-ephrinA1, and the YSA peptide. We used two fluorescent techniques, Forster Resonance Energy Transfer (FRET) and Fluorescence Intensity Fluctuations (FIF) spectrometry, to quantify the effects of the mutations on the lateral interactions between full-length EphA2 molecules in the plasma membrane.

## Materials and Methods

### Plasmid constructs

The EphA2 plasmid in the pcDNA3.1(+) vector encodes for human EphA2 tagged at the C-terminus with a fluorescent protein (either eYFP or mTurquoise) via a 15 amino acid GGS_5_ linker.(46) The glycine zipper (GZ, G540I, G544I) and heptad repeat (HR, G539I, A542I, G553I) variant were cloned using the QuikChange II Site-Directed Mutagenesis Kit according to manufacturer’s instructions (Agilent Technologies, #200523). All plasmids were sequenced to confirm the correct sequences (Genewiz).

### Cell culture and transfection

HEK293T cells were purchased from American Type Culture Collection (Manassas, VA, USA). The cells were cultured in Dulbecco’s modified eagle medium (Gibco, #31600034) supplemented with 10% fetal bovine serum (HyClone, #SH30070.03), 20 mM D-Glucose and 18 mM sodium bicarbonate at 37 °C in a 5% CO_2_ environment.

24 hours prior to transfection, cells were seeded in 35 mm glass coverslip, collagen coated Petri dishes (MatTek, P35GCOL-1.5-14-C) at a density of 2.5*10^5^ cells per dish to reach ∼70% confluency at the day of the experiment. For transfection, Lipofectamine 3000 (Invitrogen, #L3000008) was used according to manufacturer’s protocol. Single transfections were performed using 1 – 3 ug plasmid DNA. Co-transfections were performed with 1 – 4 ug total plasmid DNA in a 1:3 donor:acceptor ratio. 12 hours after transfection the cells were rinsed twice with phenol-red free, serum free starvation media and then serum starved for at least 12 hours. For experiments with added ligand, the starvation media was supplemented with 0.1% BSA to coat the wall of the dishes.

### Two photon microscopy

Before imaging, HEK293T cells were subjected to reversible osmotic stress by replacing the serum-free media with a 37°C, 1:9 serum-free media:diH_2_O, 25 mM HEPES solution. In cells the plasma membrane is normally highly ruffled and its topology in microscope images is virtually unknown.(47) The reversible osmotic stress eliminates these wrinkles and allows to convert effective 3D protein concentrations into 2D receptor concentrations.(47) In experiments with ligands, the swelling media was supplemented with 200 nM monomeric EprhinA1 (Novoprotein, #CA70), 50 nM dimeric EphrinA1-Fc (R&D Systems, #602-A1-200) or 50 μM of the engineered peptide ligand YSA (YSAYPDSVPMMSGSGSK). The cells were allowed to equilibrate for 10 minutes at room temperature. FSI-FRET, a quantitative fluorescent microscopy imaging and analysis technique, was used to measure donor (EphA2-mTurquoise) concentrations, acceptor (EphA2-eYFP) concentrations, and FRET efficiencies in micron-sized regions of the plasma membrane (47). images of cells (100 to 350 cells per condition, see Table 2) were acquired using a two-photon microscope equipped with the OptiMiS spectral imaging system (Aurora Spectral Technologies). Two scans were performed for every cell– a FRET scan (λ_1_=840 nm) in which the donor (mTurquoise) is primarily excited and an acceptor scan (λ_2_=960 nm) in which the acceptor (eYFP) is primarily excited. The output of each scan is composed of 300×440 pixels, where every pixel contains a full fluorescence spectrum in the range of 420 – 620 nm. Images of cells were acquired for up to 2 hours.

### Statistical energy test

The statistical energy test (48,49), which allows the comparison of multivariate data sets without binning, was used to compare FRET data clouds (FRET efficiencies versus concentrations of donor-labeled and acceptor-labeled proteins) of the acquired datasets.(50) Two mathematical expressions were used to calculate the statistical energy ϕ_XY_ between the two data sets X and Y, given by equations (1) and (3) below.

According to Aslan & Zech(48), ϕ_XY_ is defined as:

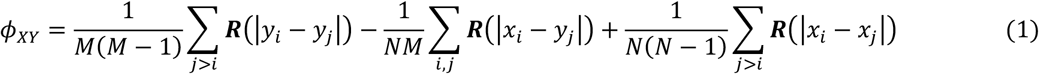

N is the number of data points in data cloud X and M is the number of data points in data cloud Y. x_i,j_ and y_i,j_ are specific data points in X and Y, respectively.

**R** in equation (1) is the logarithmic distance function:

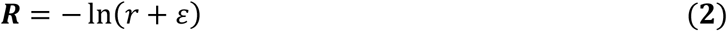

r is the Euclidean distance between the two datapoints and ε is a constant set to 1.5 to avoid singularity at r = 0.

The second way to calculate the statistical energy has been proposed by Székely & Rizzo (49) and is given by:

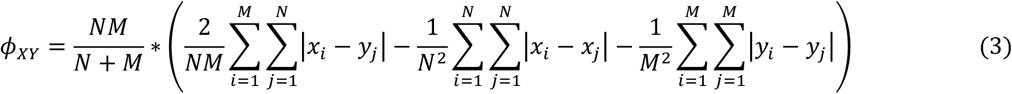

Since the test statistics are unknown, a permutation test is performed by combining the datapoints in the two data sets and generating 10^4^ sets of two new data sets with N and M randomly subsampled datapoints. Statistical energies ϕ_permutation_ for the 10^6^ random sets of binding curves were calculated to yield the test statistics. The statistical energy ϕ_XY_ for binding curves X and Y was compared to the test statistics to calculate a p-value according to

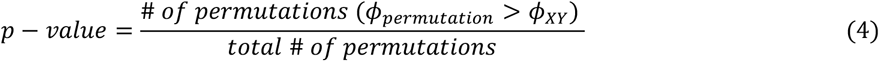

The derived p-value was adjusted with Bonferroni’s correction. A cutoff of 5% was set as the significance level.

### Fluorescence Intensity Fluctuations (FIF) Spectroscopy and analysis

FIF experiments were performed with a TCS SP8 confocal microscope (Leica Biosystems, Wetzlar, Germany) equipped with a HyD hybrid detector. Images (1024×1024, 12bit) of plasma membranes of cells expressing EphA2-eYFP were acquired in photon counting mode with a scanning speed of 20 Hz and a 488 nm diode laser excitation. The emission spectra of eYFP were collected from 520-580nm.

eYFP was excited using a 488nm diode laser at 0.1% to avoid photobleaching, at a scanning speed of 20Hz. Cells were subjected to osmotic stress with a hypoosmotic media of 75% water. This swelling minimizes the effect of ruffles, folds, invaginations, or other irregularities in the plasma membrane, while also preventing endocytosis of the receptor.

A total of ∼100 to 150 cells were imaged and analyzed. A large region in the plasma membrane was selected for each cell, and was then divided into segments of 15×15 (225 pixels^2^) as described (51), yielding a total of ∼10,000 segments per ligand. Histograms of pixel intensities were constructed for each segment, and fitted with a Gaussian function, yielding two parameters: <*I*_*segment*_>, the center of the Gaussian, and *σ*_*segment*_, the width of the Gaussian for each segment. The molecular brightness of each segment *ε*_*segment*_ was calculated as:

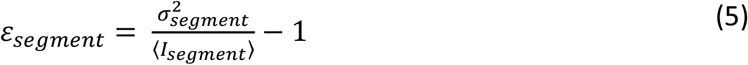

The brightness values from thousands of segments were binned and histogrammed.

These brightness distribution were fitted using Origin Lab with a log-normal function given by:

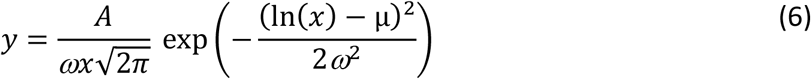

Here μ is the mean of respective ln(x) Gaussian distribution and ω is the width of the distribution. These two parameters were used to calculate the mean, median, and mode of the log-normal distribution according to:

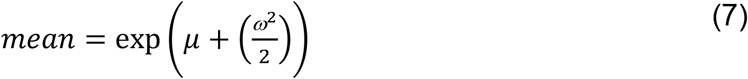

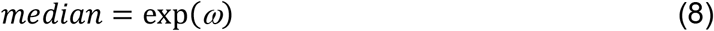

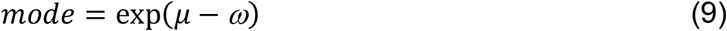

Statistical analysis was performed using ANOVA in Prism.

## Results

We characterized the homo-association of EphA2 wild-type, the G539I/A542I/G553I HR mutant, and the G540I/G544I GZ mutant in the plasma membrane of HEK293T cells using FRET. The mutations were introduced in full-length EphA2 labeled at the C-terminus with mTurquoise (mTurq, the donor) or eYFP (the acceptor), attached via a (GGS)_5_ flexible linker. We have shown that these fluorescent proteins do not perturb EphA2 autophosphorylation (52).

We used a quantitative FRET technique termed Fully Quantified Spectral Imaging FRET (FSI-FRET), which involves the acquision of complete FRET and acceptor spectra using a two-photon microscope (47). This technique employs an assumption-free, fully resolved system of equations to calculate (i) the donor concentrations, (ii) the acceptor concentrations, and (iii) the FRET efficiencies in hundreds of live cells (47). We used transient transfections to vary EphA2 expression over a broad range, from ∼100 to 10,000 receptors per square micron, and we combined data from at least 100 cells to obtain FRET binding curves.

Experiments to compare EphA2 wild-type with the HR and GZ mutants were performed in the absence of ligand as well as in the presence of saturating/near saturating concentrations of ephrinA1-Fc (50 nM), m-ephrinA1 (200 nM) or YSA peptide (50 μM) (40) to ensure that most EphA2 receptors were ligand bound, and therefore that essentially only liganded dimers/oligomers contributed to the FRET signal. Figure 1 shows the measured FRET efficiencies as a function of donor (EphA2-mTurq), acceptor (EphA2-eYFP), and total receptor concentrations, with each data point corresponding to one cell. By eye, the FRET data for the two EphA2 mutants appear very similar to the data for EphA2 wild-type. Furthermore, TM mutations cause only very modest decreases in the mean FRET intensities (Table 1). However, the mean FRET intensities do not account for differences in total expression or in donor to acceptor ratios, which are parameters that directly affect the measured FRET efficiencies for the three EphA2 variants. Thus, the values in Table 1 cannot be directly compared in a meaningful way.

**Figure 1.**
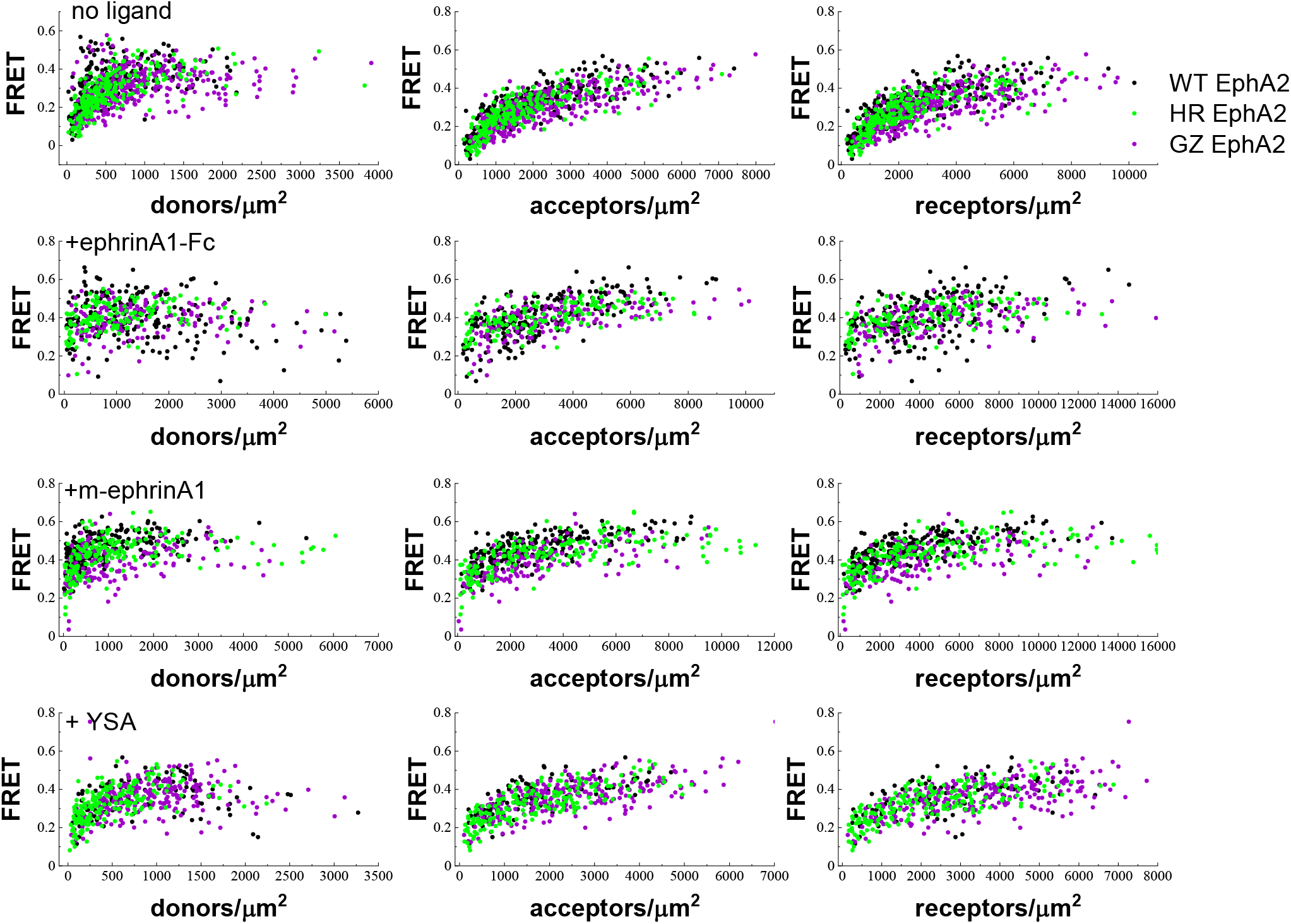
FRET data for EphA2 (black), the EphA2 GZ mutant (purple), and the EphA2 HR mutant (green). Row 1 shows the data with no added ligand, row 2 with 50 nM ephrinA1-Fc, row 3 with 200 nM m-ephrinA1, and row 4 with 50 μM YSA. Column 1 shows FRET as a function of donor concentration, column 2 as a function of acceptor concentration, and column 3 as a function of total receptor concentration.

**Table 1.** Number of cells analyzed in the FRET experiments, mean FRET intensities and standard errors. FRET efficiencies are averaged over all EphA2 concentrations. The mutations either decrease the average FRET, or have no measurable effects.

|  |  | **FRET** | **N** |
| --- | --- | --- | --- |
| **no ligand** | WT | $0.32 \pm 0.01$ | 255 |
| | HR | $0.28 \pm 0.01$ | 236 |
| | GZ | $0.30 \pm 0.01$ | 359 |
| **ephrinA1-Fc** | WT | $0.40 \pm 0.01$ | 196 |
| | HR | $0.40 \pm 0.01$ | 144 |
| | GZ | $0.39 \pm 0.01$ | 156 |
| **m-ephrinA1** | WT | $0.46 \pm 0.01$ | 226 |
| | HR | $0.42 \pm 0.01$ | 187 |
| | GZ | $0.39 \pm 0.01$ | 152 |
| **YSA** | WT | $0.36 \pm 0.01$ | 105 |
| | HR | $0.33 \pm 0.01$ | 193 |
| | GZ | $0.36 \pm 0.01$ | 257 |

Thus, we sought to determine if there are statistically significant differences in the measured FRET efficiencies as a function of donor and acceptor concentrations due to the mutations. However, comparison of the FRET data clouds for EphA2 wild-type and mutants (Figure 1) is not trivial, because FRET values depend on two parameters (EphA2-mTurq concentration and EphA2-eYFP concentration) that vary independently in each cell. Thus, we reasoned that we needed a new methodology that allows analysis of multidimensional data in order to determine the statistical significance of differences between data clouds such as those shown in Figure 1.

We applied a statistical test known as ‘statistical energy’, which meets the above criteria (48,49). This test has been mainly used for particle physics applications. Here we use it, for the first time, to compare protein interaction data. In the statistical energy test, each datapoint is viewed as an electrical charge, and different FRET clouds (FRET efficiencies as a function of donor and acceptor concentrations) are assumed to have different charge signs. Therefore, if two FRET clouds are very different, statistical energy is considered high due to charge repulsion. However, if two FRET clouds overlap extensively, statistical energy will be low. Two different forms of the statistical energy, ϕ_XY_ (where X and Y refer to the two data sets being compared), can be calculated (according to equations (1) and (3) in the Materials and Methods) and used for statistical analysis. To determine p-values non-parametrically, a numerical permutation test was performed by creating 10,000 random permutations of each FRET cloud. The associated statistical energy distributions were used to yield the test statistics according to equation (4). These statistical analyses show that both the HR and the GZ mutations have a significant effect on FRET efficiencies (compared to EphA2 wild-type) in the cases of no ligand, ephrinA1-Fc and m-ephrinA1 (Table 2). In the presence of YSA, the HR and GZ mutations have no effect. The FRET efficiencies of the two mutants are not significantly different from each other in the absence of ligand and in the presence of ephrinA1-Fc, but are significantly different in the presence of m-ephrinA1 and YSA. The two different forms of the statistical energy, ϕ_XY_, yield consistent statistical significance for all comparisons (Table 1), supporting the validity of the results.

**Table 2.** Statistical analysis of the EphA2 FRET datasets. The p-values were calculated with a statistical energy test, where the statistical energies are given by either equation (1) or equation (3) (energy definitions according to Aslan and Zech (48) and Székely and Rizzo (49), respectively). The p-values for the two energy definitions are in good agreement. Both TM helix mutations have a significant effects on the FRET efficiencies in the cases of no ligand, ephrinA1-Fc and m-ephrinA1. In the presence of YSA the HR mutation has a marginally significant effect, while the GZ mutation has no effect. The two mutations have similar FRET efficiencies in the absence of ligand and in the presence of ephrinA1-Fc, but significantly different FRET efficiencies in the presence of m-ephrinA1 and YSA.

| | | **p-value**<br>$\phi_{XY}$ (eq. (1)) | **p-value**<br>$\phi_{XY}$ (eq. (3)) |
| --- | --- | --- | --- |
| **no ligand** | WT vs GZ | 0.0054 | 0.0033 |
|  | WT vs HR | 0.0003 | 0.0003 |
|  | GZ vs HR | >0.05 | >0.5 |
| **ephrinA1-Fc** | WT vs GZ | 0.0087 | 0.0111 |
|  | WT vs HR | 0.0039 | 0.0087 |
|  | GZ vs HR | >0.05 | >0.05 |
| **m-ephrinA1** | WT vs GZ | 0.0003 | 0.0003 |
|  | WT vs HR | 0.0006 | 0.0003 |
|  | GZ vs HR | 0.0027 | 0.0012 |
| **YSA** | WT vs GZ | >0.05 | >0.5 |
|  | WT vs HR | 0.0522 | 0.0528 |
|  | GZ vs HR | 0.0006 | 0.0015 |

The mechanistic interpretation of the FRET data is not straightforward, because FRET depends not only on dimerization/oligomerization strengths, but also on conformational effects that may affect the positioning and dynamics of the fluorescent proteins in the oligomers (53,54). Furthermore, FRET cannot differentiate between high order oligomers of different sizes or provide information on the heterogeneity in the populations of oligomers (54). To gain further insight into the effects of the HR and GZ TM mutations on EphA2 dimerization/oligomerization, we used fluorescence intensity fluctuations (FIF) spectrometry. FIF measures molecular brightness, which is known to scale with oligomer size (51). For the FIF experiments, HEK293T cells were transiently transfected with plasmids encoding wild-type and mutant EphA2-eYFP as well as LAT-eYFP as a monomer control (55,56) and E-cadherin-eYFP as a dimer control (57). Following expression of the fluorescent proteins, the plasma membrane in contact with the substrate was imaged in a confocal microscope and molecular brightness values were calculated in small regions of the plasma membrane (15×15 pixels) according to equation (5), as described (51).

In the absence of ligand, the brightness distributions for EphA2 wild-type and both the HR and GZ mutants are between the distributions of LAT and E-cadherin, indicating a mixture of EphA2 monomers and dimers (Figure 2). In the presence of ephrinA1-Fc and m-ephrinA1, EphA2 wild-type and mutant brightness distributions are shifted to higher brightness compared to E-cadherin, indicating the formation higher order oligomers in addition to dimers. In contrast, in the presence of the YSA peptide, the brightness distributions for EphA2 wild-type and both mutants are similar to the E-cadherin distribution, indicating that the peptide ligand promotes EphA2 dimerization, as previously proposed based on FRET data (58).

**Figure 2.**
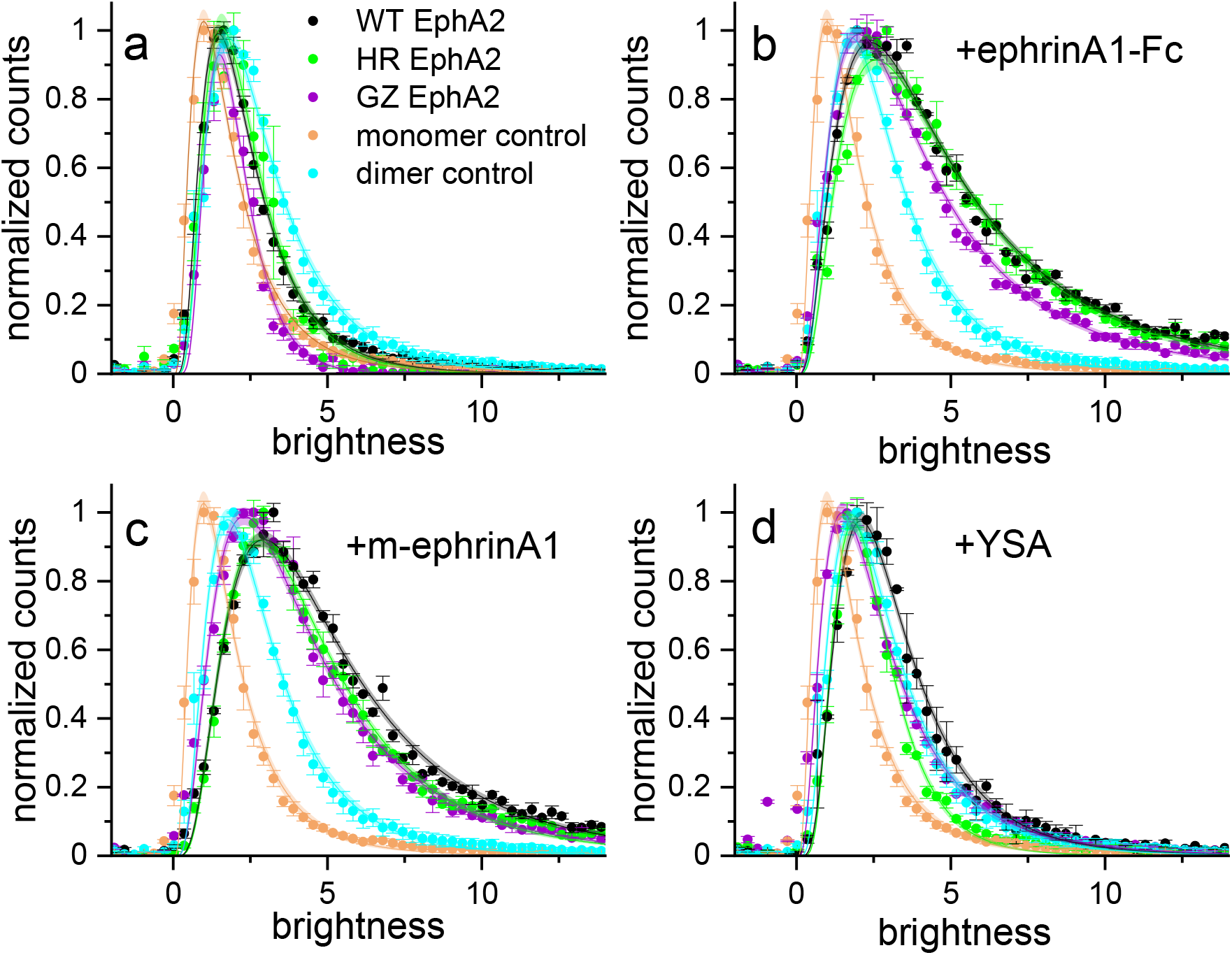
FRET data for EphA2 (black), the EphA2 GZ mutant (purple), and the EphA2 HR mutant (green).The molecular brightness distributions for EphA2 with no added ligand (a), 50 nM ephrinA1-Fc (b), 200 nM m-ephrinA1 (c), and 50 μM YSA (d). The monomer control LAT (orange) and the dimer control E-cadherin (turquoise) are shown in each panel.

The FIF distributions are well described by a log-normal distribution, which is characterized by two best-fit parameters (µ and ω, see equation (6)). These two parameters are used to calculate the median, mean and mode of the log-normal distributions (Table 3), enabling statistical comparisons of the FIF data to determine if there are significant differences in the oligomer size. Comparing the means of the FIF brightness distributions show that the GZ mutation has a smaller oligomer size with a statistically significant difference from EphA2 wild-type and **the** HR **mutant**, both in the absence of ligand and in the presence of ephrinA1-Fc (Table 4). Statistical analyses comparing the means of the FIF brightness distributions for EphA2 wild-type and the two TM mutants show that only the GZ mutation has a statistically significant effect in the absence of ligand and in the presence of ephrinA1-Fc (Table 4). In contrast, both HR and GZ mutations significantly shift the FIF distributions to lower brightness values, compared to EphA2 wild-type, in the presence of m-ephrinA1 and YSA (Tables 3 and 4). When comparing the two mutants to each other, the differences are statistically significant in the cases of no ligand, ephrinA1-Fc and YSA, but not in the case of m-ephrinA1. A comparison of the mutants to each other shows that the HR mutant exhibits higher mean brightness without ligand, with m-ephrinA1 and with ephrinA1-Fc. However, in the presence of YSA, the HR mutant has lower mean brightness than the GZ mutant.

**Table 3.**
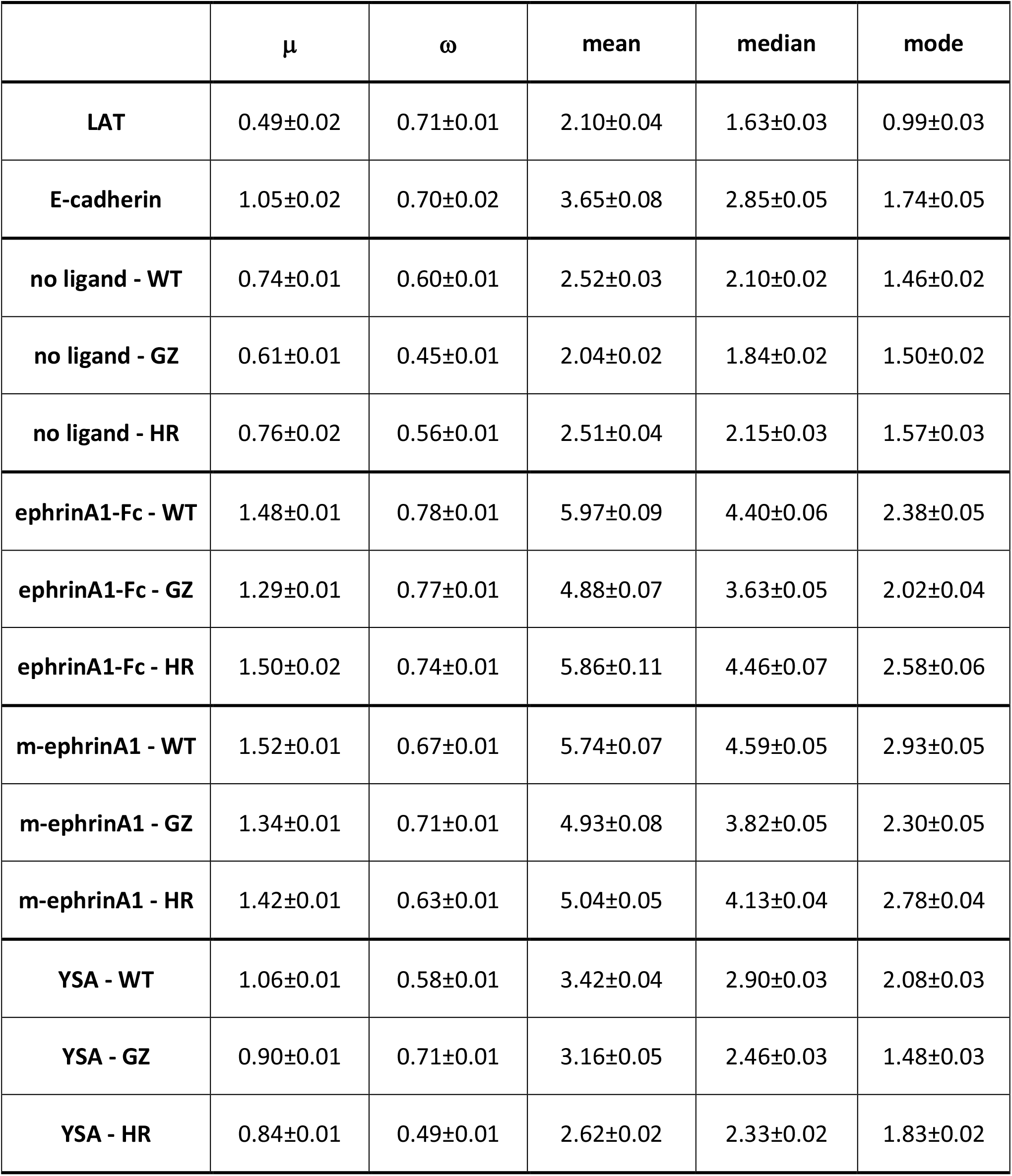
Best-fit parameters for the FIF distributions. µ and ω are the two best-fit parameters of the log-normal distribution (see equation (6)). These two parameters are used to calculate the median, mean and mode of the log-normal distribution according to equations (7), (8) and (9), respectively.

| | $\mu$ | $\omega$ | mean | median | mode |
| --- | --- | --- | --- | --- | --- |
| **LAT** | 0.49±0.02 | 0.71±0.01 | 2.10±0.04 | 1.63±0.03 | 0.99±0.03 |
| **E-cadherin** | 1.05±0.02 | 0.70±0.02 | 3.65±0.08 | 2.85±0.05 | 1.74±0.05 |
| **no ligand - WT** | 0.74±0.01 | 0.60±0.01 | 2.52±0.03 | 2.10±0.02 | 1.46±0.02 |
| **no ligand - GZ** | 0.61±0.01 | 0.45±0.01 | 2.04±0.02 | 1.84±0.02 | 1.50±0.02 |
| **no ligand - HR** | 0.76±0.02 | 0.56±0.01 | 2.51±0.04 | 2.15±0.03 | 1.57±0.03 |
| **ephrinA1-Fc - WT** | 1.48±0.01 | 0.78±0.01 | 5.97±0.09 | 4.40±0.06 | 2.38±0.05 |
| **ephrinA1-Fc - GZ** | 1.29±0.01 | 0.77±0.01 | 4.88±0.07 | 3.63±0.05 | 2.02±0.04 |
| **ephrinA1-Fc - HR** | 1.50±0.02 | 0.74±0.01 | 5.86±0.11 | 4.46±0.07 | 2.58±0.06 |
| **m-ephrinA1 - WT** | 1.52±0.01 | 0.67±0.01 | 5.74±0.07 | 4.59±0.05 | 2.93±0.05 |
| **m-ephrinA1 - GZ** | 1.34±0.01 | 0.71±0.01 | 4.93±0.08 | 3.82±0.05 | 2.30±0.05 |
| **m-ephrinA1 - HR** | 1.42±0.01 | 0.63±0.01 | 5.04±0.05 | 4.13±0.04 | 2.78±0.04 |
| **YSA - WT** | 1.06±0.01 | 0.58±0.01 | 3.42±0.04 | 2.90±0.03 | 2.08±0.03 |
| **YSA - GZ** | 0.90±0.01 | 0.71±0.01 | 3.16±0.05 | 2.46±0.03 | 1.48±0.03 |
| **YSA - HR** | 0.84±0.01 | 0.49±0.01 | 2.62±0.02 | 2.33±0.02 | 1.83±0.02 |

**Table 4.** P values for comparisons of FIF brightness distributions. P < 0.05 indicates statistical significance.

|  | **p value** |
| --- | --- |
| **no ligand** |  |
| WT vs HR | >0.05 |
| WT vs GZ | <0.0001 |
| HR vs GZ | <0.0001 |
| **ephrinA1-Fc** |  |
| WT vs HR | >0.05 |
| WT vs GZ | <0.0001 |
| HR vs GZ | <0.0001 |
| **m-ephrinA1** |  |
| WT vs HR | <0.0001 |
| WT vs GZ | <0.0001 |
| HR vs GZ | >0.05 |
| **YSA** |  |
| WT vs HR | <0.0001 |
| WT vs GZ | <0.0001 |
| HR vs GZ | <0.0001 |

## Discussion

Despite many years of RTK research, the physiological role of the TM helix in receptor dimerization/oligomerization and signaling remains poorly understood (59). In part, this is because much of the knowledge about RTK structure-function relationships has come from X-ray and NMR-derived structures of isolated RTK domains. For instance, there are crystal structures of the isolated EphA2 extracellular region, kinase domain, and intracellular region as well as NMR structures of the isolated EphA2 TM helix (20,31-33). Such structures cannot reveal how ligand-induced structural changes in the extracellular region are relayed to the intracellular region through the TM helix. Thus, mutagenesis guided by structural studies is often used to gain insight into the role of the different parts of an RTK in activation and signaling.

Since RTKs function via oligomerization, amino acid residues that mediate intermolecular contacts in crystals are potentially important for RTK function. Crystals of the EphA2 extracellular region are stabilized via two distinct interfaces, the “dimerization” and the “clustering” interface (31,32). The “dimerization” interface involves both receptor-receptor and receptor-ephrin contacts in the ligand-binding domain, including contacts involving amino acid G131. The “clustering” interface involves contacts between residues L223, L254 and V255 in the EphA2 cysteine-rich region, without ligand participation. In prior studies involving mutagenesis and FRET measurements, we have shown that the G131Y mutation and/or the triple L223R/L254R/V255R mutation reduce the stabilities of EphA2 full-length dimers and oligomers (with dissociation constants increasing up to 5-10-fold as a consequence of the mutations) (45). Furthermore, the two sets of destabilizing mutations have been shown to substantially impair cross-phosphorylation of tyrosine residues in EphA2 as well as the migration of EphA2-expressing cells (31,32,45). Therefore, the crystal structures of the EphA2 extracellular region successfully capture biologically relevant EphA2 interactions.

Here we used mutagenesis to study the effects of EphA2 TM helix interface modifications on EphA2 interactions in the plasma membrane. We used two fluorescence techniques, FRET and FIF, to assess EphA2 dimerization/oligomerization. These two techniques yield different information because FRET efficiencies depend on both the interaction propensities and the distance between fluorescent proteins, where the distance depends on the structure of the EphA2 dimers and oligomers. On the other hand, FIF is not affected by structural parameters. We mutated residues previously reported to mediate contacts between the isolated TM helices based on an NMR structure, as well as contacts identified in isolated TM helices in the plasma membrane of cells (19,24). The two techniques yielded similar results for some receptor-ligand combinations but not others, consistent with the fact that FRET and FIF measure different parameters. We found that destabilization of the two TM interfaces as a result of the mutations only modestly affects the measured FRET efficiencies and FIF brightness distributions, in contrast to the substantial effects of the extracellular interface mutations. This suggests that the interactions between the TM helices either lack strong sequence specificity or do not strongly contribute to receptor-receptor association. Therefore, the nature and importance of the interactions between the extracellular regions and between the TM helices within full-length EphA2 appear fundamentally different.

It has long been believed that in RTKs one TM helix dimer conformation corresponds to one specific dimeric conformation of the extracellular region (13,60,61). For EphA2, the TM helix has been proposed to associate through the HR interface in the inactive state and to switch to the GZ interface in the active state (24). We reasoned that if this were the case, the HR and GZ TM helix mutations should have distinctly different effects on the measured FRET efficiencies and FIF distributions in the absence of ligand and in the presence of ligands that essentially lack agonistic ability (such as the YSA peptide) compared to ligands that strongly activate EphA2 (ephrinA1-Fc and m-ephrinA1). Our data do not confirm this expectation. In the FRET experiments, we found that both HR and GZ mutations alter the FRET signal both in the absence of ligand (inactive EphA2) and in the presence of ephrinA1-Fc and m-ephrinA1 (active EphA2). Similarly, in the FIF experiments we observed that both HR and GZ mutations decrease the means of the brightness distributions in the presence of m-ephrinA1 (active EphA2) and in the presence of the YSA peptide (inactive EphA2). In addition, the GZ mutation, but not the HR mutation, decreases the FIF mean brightness distributions both in the absence of ligand (inactive EphA2) and in the presence of ephrinA1-Fc (active EphA2). Thus, our data do not support the current model of EphA2 activation via a TM dimer switch because the two interfaces appear to simultaneously play a role in both active and inactive forms of EphA2 and in both unliganded and liganded EphA2. Notably, other functions have been proposed for the EphA2 TM helix, for example in the regulation of EphA2 localization at epithelial cell-cell junctions (62).

EphA2 is known to form both dimers and oligomers, depending on the identity of the bound ligand (45). Here we find that the HR and GZ mutations do not fundamentally change the oligomerization state of EphA2, but rather cause only modest shifts in dimer/oligomer populations in some cases. EphA2 wild-type, as well as the HR and GZ mutants, are all dimeric both in the absence of ligand and in the presence of the YSA peptide. The mutations tend to modestly decrease the average brightness in the FIF experiments, which likely means that the unliganded and YSA-bound EphA2 dimers are destabilized by both TM helix mutations leading to enrichment of the monomer populations. Binding of ephrinA1-Fc or m-ephrinA1 leads to the appearance of higher older oligomers for all EphA2 mutants. In previous work, we concluded on the basis of FRET data that EphA2 in the presence of m-ephrinA1 is best described by a dimer model (45). The FIF experiments, however, reveal a heterogeneous population of dimers and oligomers of different sizes, which cannot be distinguished in the FRET experiments (54). Both TM mutations shift EphA2 oligomer size distributions to lower values in the presence of ephrinA1-Fc and m-ephrinA1. Notably, in both FRET and FIF experiments the effects of the two mutations (HR versus GZ comparison in Tables 2 and 4) are different for ephrinA1-Fc and m-ephrinA1, two ligands that can differentially bias EphA2 signaling responses (40). This suggests that the EphA2 molecules may have different arrangements in the ephrinA1-Fc- and m-ephrinA1-bound oligomers, which may contribute to ligand bias.

The effects of the HR and GZ TM helix mutations on EphA2 phosphorylation were also found to be relatively modest in previous studies (19). Mutations in the HR motif increased EphA2 phosphorylation, while mutations in the GZ motif decreased EphA2 phosphorylation compared to EphA2 wild-type (19). These effects were observed both in the absence of ligand and in the presence of ephrinA1-Fc. Interestingly, here we find that the GZ mutation also decreases the mean of the brightness distribution both in the absence of ligand and in the presence of ephrinA1-Fc, but the HR mutation does not. Thus, the phosphorylation decrease due to the GZ mutation observed in prior work (24) may be related to a small decrease in average EphA2 oligomer size, inferred from the decrease of the mean brightness observed here.

It has been proposed that the GZ interface is important for EphA2 function (24). Our findings are generally consistent with this view, since in FIF experiments destabilization of the GZ interface reduces the oligomer size in all cases. However, the alternate HR interface also affects the FIF distributions in the cases of m-ephrinA1 and YSA. Thus, it appears that both interfaces play a role in the stabilization of the m-ephrinA1-bound oligomers and YSA-bound dimers. It is not clear if the TM helices in the ephrinA1-Fc and m-ephrinA1-stabilized oligomers interact exclusively as dimers, or whether a TM helix can interact with more than one other TM helix forming a trimer. It is also possible that EphA2 dimer/oligomer conformations exist where the TM helices are not in direct contact, perhaps providing an explanation for the small effects of the mutations. While many questions remain unanswered, one way to interpret our results is to hypothesize that the TM helices in EphA2 dimers and oligomers can interact promiscuously and transiently, via different non-unique interfaces. The existence of an ensemble of TM helix conformations populated within an RTK dimer or oligomer could explain why RTK TM helices have remained unresolved in cryo-EM structures.

A question arises as to whether an ensemble of TM helix conformations can have functional implications in EphA2 signaling, given that the EphA2 juxtamembrane segment is flexible. It can be speculated that plasticity in TM helix contacts may augment the overall conformational space that is accessible to the EphA2 intracellular region. This could facilitate the formation of diverse intracellular arrangements of EphA2 dimers and oligomers to ensure their ability to cross-phosphorylate and/or to phosphorylate substrate proteins. We propose that this structural plasticity could be critical for the ability of EphA2 to trigger a multitude of downstream signaling cascades.

## Funding

Supported by NIH grants R01GM131374 (EBP and KH), R01GM068619 (KH), and NSF MCB 2106031 (KH).

## Notes

### Competing Interest Statement

The authors have declared no competing interest.

